# When a rose is not a rose: Pyramidal neurons respond differently in healthy and epileptic human neocortex

**DOI:** 10.1101/185710

**Authors:** Kevin Joseph, Vidhya M. Ravi, Katerina Argiti, Thomas J. Feuerstein, Josef Zentner, Rosanna Migliore, Michele Migliore, Ulrich G. Hofmann

## Abstract

Epilepsy affects a huge number of patients by severe disruption of brain functions and is characterised by recurrent seizures, sometimes hard to be treated by medications. Seizure induced cellular consequences in ionic gradient and homeostasis are expected to result in electrophysiological differences between epileptic and non-epileptic neurons.

In the following work we demonstrate these differences in layer III cortical pyramidal neurons sourced from epileptic and non-epileptic human patient tissue. Although visually indistinguishable and featuring similar membrane potentials and latency to first spikes upon whole cell patch stimulation, epileptic pyramidal neurons display a larger rheobase and a smaller membrane resistance, responding less efficiently to electrical stimulation than their peri-tumorous equivalents. This decreased excitability contradicts results in comparable animal models of epilepsy and was further corroborated by detailed analysis of spiking characteristics and phase plot analysis of these events. Both point to an overexpression of K^+^ channels trying to compensate for the hyperexcited, epileptic network state. A computational model of a pyramidal neuron was utilized to give an estimate of the needful relative changes in K^+^ and Na^+^ conductances.

## 1. Introduction

Epilepsy is a surprisingly common debilitating condition characterized by recurrent seizures. There is no clear understanding to why seizures occur based on our current knowledge from neuro(patho)physiology or experimental studies. This is due to the fact that epilepsy involves numerous factors such as genetic, cellular-level, excitatory/inhibitory behaviour, changes in ionic conductances, number of synaptic connections etc. (Chakravarthy et al., 2009). An epileptic seizure entailing paroxysmal sodium influx into the cells involves increased excitation along with a hyper-synchronous state, which leads to a temporary breakdown of homeostatic control due to the intense ion flux (Badawy et al., 2012; Lindsay, 2014). Mechanisms involved in the necessary intra- and extracellular regulation of ionic concentrations are hard to investigate; however, it has been demonstrated that neuronal activities are intrinsically modulated by the ionic gradients across their cellular membranes, which depend on the complex interaction of mechanisms related to ionic homeostatic regulation (Miranda et al., 2013). Homeostatic adaptability is key to maintaining functional stability, while providing room for developmental changes (Ciarleglio et al., 2015). This has been shown to coordinate changes of different cellular properties, like changes in intrinsic excitability due to increased excitatory synaptic drive (Debanne et al., 2003; Desai, 2003; O’Leary et al., 2013; Turrigiano et al., 1994). Evidence has been presented that the presence of any activity, like adapting to homeostatic challenges, modulates intrinsic neuronal excitability (Franklin et al., 1992; Moody, 1998; Turrigiano et al., 1995).

Since pyramidal cells are the primary excitatory cells in the neocortex and presumably most affected by any homeostatic fluctuations, they are the target of our study. The following work demonstrates compelling evidence that cortical pyramidal cells in epileptic tissue do undergo specific changes to their electrophysiological signature and excitability. Since preventing cortical neurons from firing dramatically increased their excitability mediated by selective regulation of sodium and potassium currents in homeostatic adaptation (Desai et al., 1999), one may assume a corresponding decrease in cellular excitability in epilepsy, due to the persistent occurrence of seizure events. However, this effect was not found in animal models of epilepsy, yet. To the best of our knowledge, the present study is the first experimental body of evidence trying to prove an epilepsy induced decrease in excitability.

In order to estimate the changes in intrinsic properties, computational modelling was employed as well to illuminate the role of the sodium (Na^+^) and potassium currents (K^+^) in the regulation of the excitability and intrinsic properties. We are convinced that this observation may build a step on the long journey of epileptic therapeutics for patients with pharmacoresistant epilepsy.

## 2. Results and discussion

To compare the electrophysiological response of pyramidal cells between epileptic and non-epileptic tissue, whole cell patch clamp recordings were carried out. Electrophysiological recordings were carried out from n=19 (N=4) neurons from surgically resected epileptic foci and n=14 (N=7) neurons from tumor access tissue, that was not macroscopically infiltrated with tumor.

### 2.1 Electrophysiological Characteristics

A significantly lower number of spikes evoked by stepwise depolarization of the soma of epileptic cells in comparison to non-epileptic cells. There is a significant difference between the spikes evoked at 500 pA, which is shown in (p=0.0092) (Fig. 1 a(inset i)). The rheobase was significantly higher in the case of epileptic pyramidal cells vs. non-epileptic pyramidal cells (p=0.0002) (Fig. 1 d). However, this increase in the rheobase did not affect the membrane potential at which the first action potential is observed (p=0.5591), nor the latency to the first spike (p=0.2354) (Fig. 1 b, c.). This observation can be explained by (Frankenhaeuser, 1965) who showed that a change in membrane capacitance leads to a corresponding increase in the rheobase, without a significant change in the threshold potential, as we have measured (Fig. 1 b).

**Figure 1:**
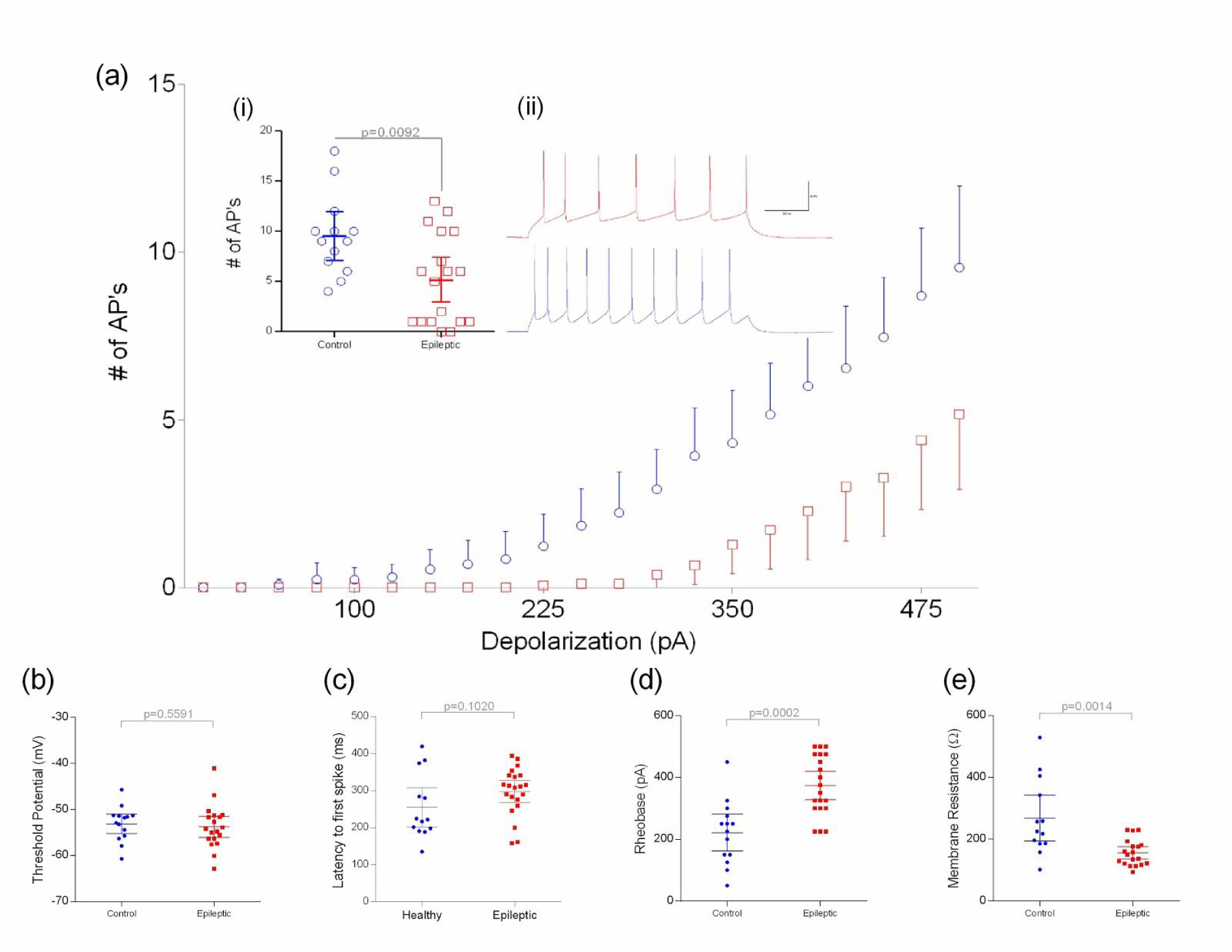
Data from non-epileptic pyramidal cells presented in blue and data from epileptic pyramidal cells presented in red (a) Number of Action potentials evoked by stepwise depolarization. **Inset (i)** Comparison of the number of action potentials evoked by 500 pA stimulation. A significantly higher number of Action potentials are observed in the case of non-epileptic tissue (p=0.0092) **Inset (ii)** Representation of the evoked action potentials to 1 second depolarization pulses at 500 pA (b) The membrane potential at which the first spikes are evoked is not significantly different between epileptic (-53.74± 4.785) and non-epileptic tissue (-53.10±3.752). (c) The same is observed for the latency to first spike between non-epileptic pyramidal cells vs epileptic pyramidal cells. (e) Stepwise depolarization experiments show that the rheobase is significantly different between non-epileptic pyramidal cells and epileptic pyramidal cells; p=0.0002. (f) The difference in the rheobase can be shown to be due to the difference in the membrane resistance between non-epileptic and epileptic pyramidal cells. All data represented as Mean ± 95% confidence intervals. Non-parametric Mann-Whitney test used for analysis.

Reduced membrane resistance (Fig. 1 e) observed in epileptic cells (78.87 ± 10.36 MΩ for non-epileptic cells vs 53.13 ± 12.17 MΩ for epileptic cells, p=0.0007), could help explain why we see such a significant difference in the rheobase. This decreased membrane resistance could be a result of increased channel activation.

**Figure 2:**
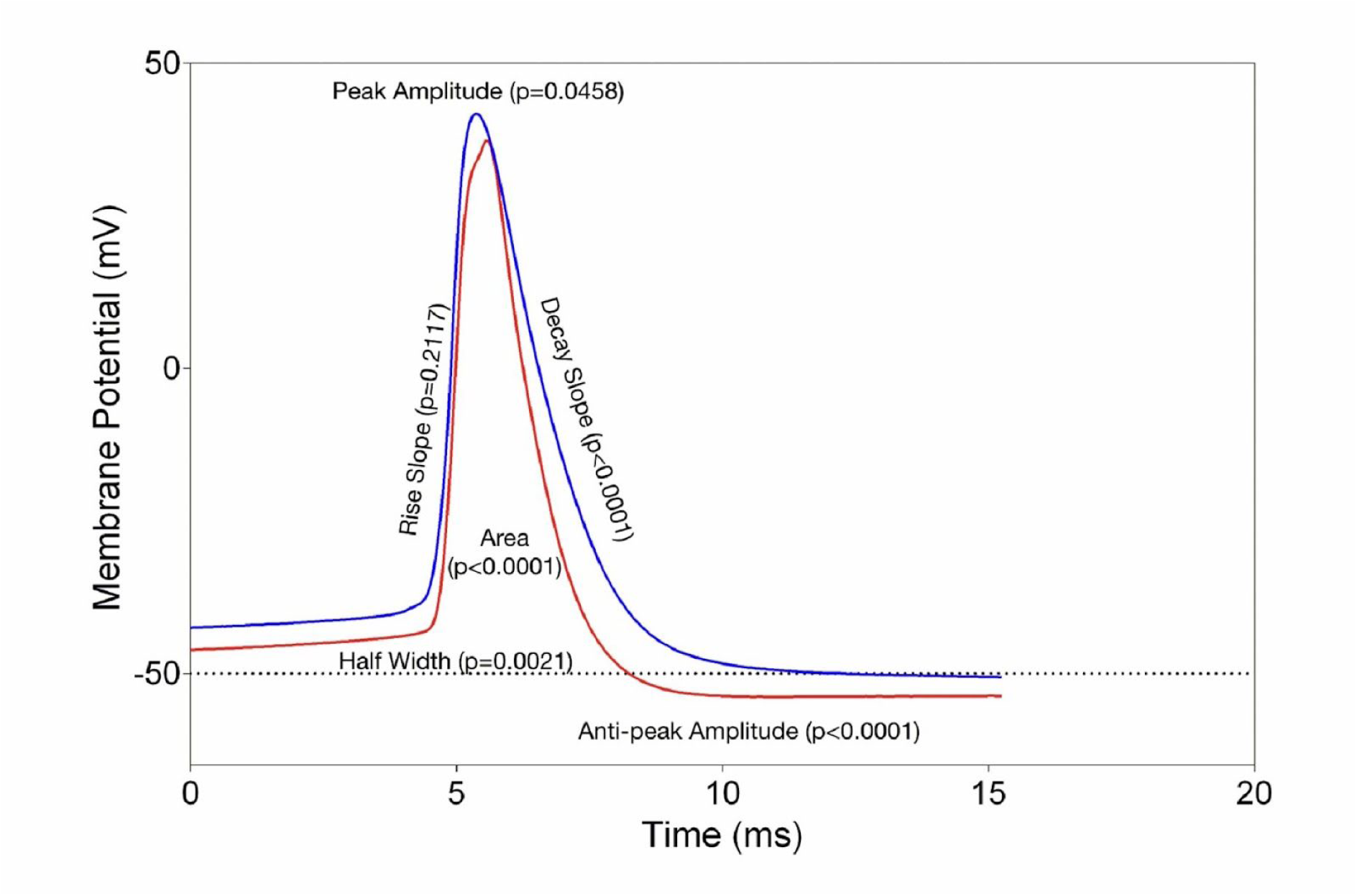
(a) Average action potential representation for non-epileptic vs epileptic pyramidal cells. non-epileptic trace represented in blue and epileptic trace represented in red. The peak amplitude, antipeak amplitude, area, half width and decay slope are all significantly different between epileptic and non-epileptic pyramidal cells. The differences are represented as p values along with the phase they represent. MW test used.

Evoked action potentials show that the Na^+^ channel activation remains similar (Rise slope, Fig. 3c), while the voltage gated K^+^ channels are activated sooner, with higher intensity (Decay slope, Fig. 3d; Anti peak amplitude, Fig. 3b). These voltage gated K^+^ currents have been proposed to counter membrane depolarization, which in turn restricts neuron excitability (Sarria et al., 2012; Vydyanathan et al., 2005). This indicates an overexpression of K^+^ channels to try and compensate for the hyperexcited network state. A significant increase in the width of the action potential (p=0.0021, Fig. 3f) could be explained by increased activation of other K^+^ currents: e.g. a delayed rectifier current and a calcium activated K^+^ current (Bean, 2007).

**Figure 3:**
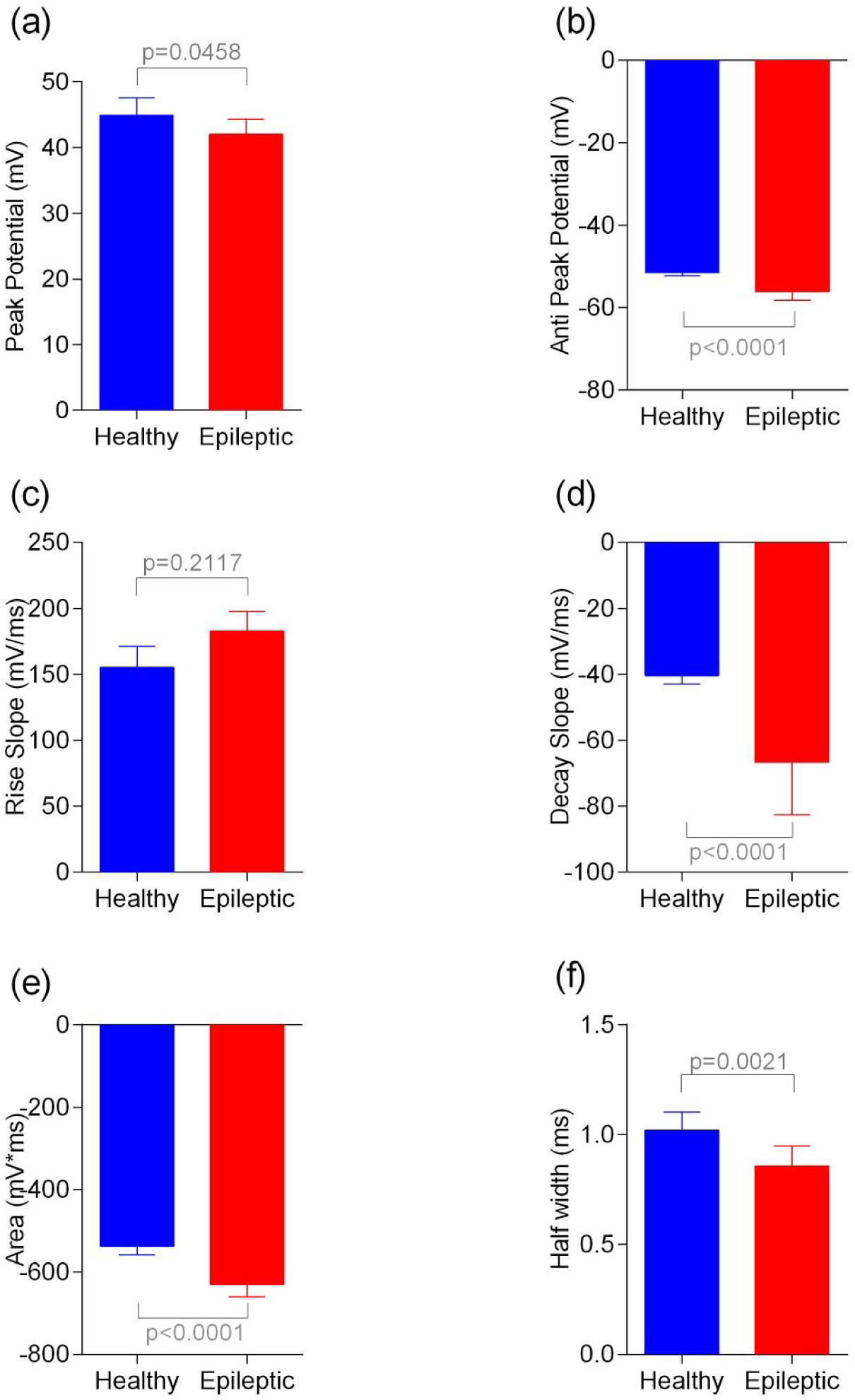
(a) There is a significant difference between epileptic cells and non-epileptic cells when it comes to the peak membrane potential (p=0.0458). (b) We see a strong difference in the anti-peak amplitude (p<0.0001) as well (c) The rise slope, which shows the Na^+^ channel active phase does not show a significant difference (p=0.2117) (d) The decay slope, which shows the K^+^ channel activity phase shows a strong difference (p<0.0001). (e) We also report a strong increase in the area action potentials in epileptic cells in comparison to non-epileptic cells (p<0.0001). (f) There is also a corresponding increase in the half-width of the action potential (p=0.0021)

### 2.2 Neuron dynamics and channel conductances

Phase plots are an eloquent way to represent changes in channel distribution and neuronal dynamics (Harnett et al., 2015). To explain found differences in the active properties of neurons from non-epileptic and epileptic patients, we produced a phase-plot (dV/dt vs. V) of the somatic membrane potential during 1 sec of suprathreshold current injections. To better isolate single spike dynamics, only current injections eliciting four action potentials were used in the analysis thus permitting sufficient time between spikes to achieve a complete and stable membrane repolarization (Fig. 4a). Only the negative part of the phase plot was significantly different between the two cases (Mann-Whitney rank sum Test, p=0.005). This region and negative derivative correspond to the repolarization phase of the action potential (Bean, 2007). Typical results from individual cells are shown in Fig. 4b, where we report the phase plot and the relative membrane potential. By looking at their features we can infer some differences between the intrinsic electrophysiological properties between neurons from non-epileptic (Fig. 4, blue) or epileptic patients (Fig. 4, red).

**Figure 4:**
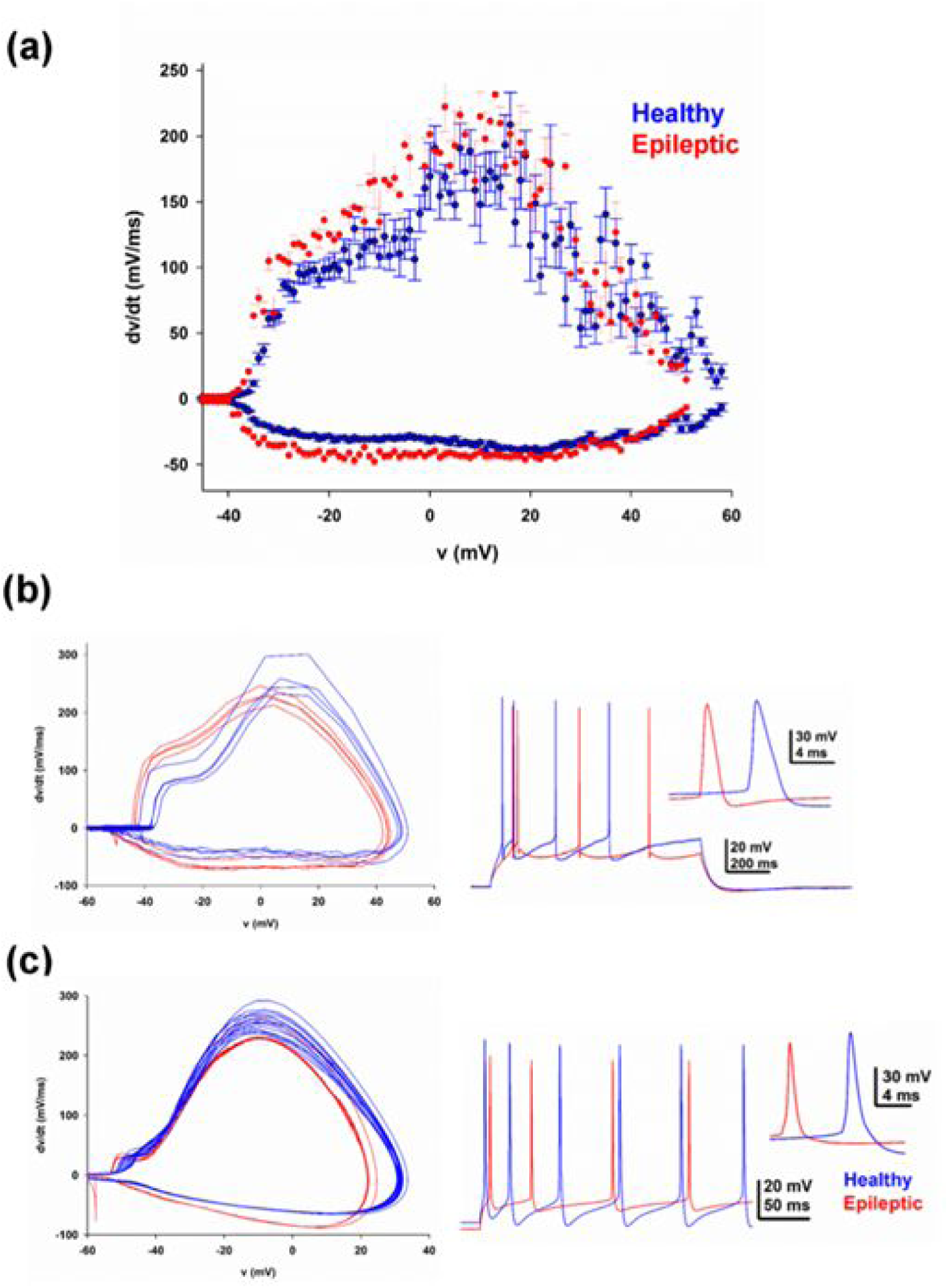
Phase-plots and typical traces from experiments and model; in all cases, blue and red traces and symbols refer to recordings of neurons from non-epileptic or epileptic patients, respectively. (a) Average phase plots of the somatic membrane potential during 2 sec somatic current injection steps eliciting four action potentials. (b) Typical phase plots from experimental recordings of individual cells (left), and the corresponding somatic membrane potential of a neuron, from a non-epileptic or an epileptic patient, in response to a depolarizing current of 500 pA or 450 pA, respectively (right). (c) Phase-plots from the model neuron (left), and the corresponding somatic membrane potential from a simulation modeling a non-epileptic or an epileptic cell, in response to a depolarizing current of 350 pA or 450 pA, respectively (right).

Two differences are clearly evident; the cell from an epileptic patient exhibited: 1) a phase plot with a larger (negative) derivative and 2) a shorter AP duration. Using a biophysically accurate model of a pyramidal neuron ((Ascoli et al., 2010; Migliore and Migliore, 2012)), we found that both features can be reproduced with a substantial increase (+50%) in the peak conductance of A-type potassium current, and compensatory changes in the M-type potassium (-90%) and the Na^+^ currents (+9%), respectively. It should be noted that each current has a role in modulating the shape of the somatic membrane potential and of its phase plot. More specifically, the increase in the K_A_ current (which is fast and transient, (Migliore et al., 1999)) can take into account both the shorter duration of an action potential and the faster repolarization phase observed experimentally. However, this increase alone would also severely reduce a cell’s excitability, and the model predicts that it must be compensated by a reduction in another potassium current (K_M_ in our case) and by a slight increase of the Na^+^ current.

## 3. Methods

### 3.1 Tissue Source

Fresh human neocortical tissue from temporal areas was obtained during therapeutic removal of epileptogenic foci (temporal lobe epilepsies, 5 patients) or subcortical brain tumors (7 patients). Tissue derived from brain tumor patients is referred to as ‘control’ group in the following text. Tissue macroscopically infiltrated with tumor was discarded. Written consent, according to the Declaration of Helsinki, as requested by the local Ethics Committee (No 187/04), was obtained for each patient. The procedure follows those described previously by (Rassner et al. 2016). The tissue derived from N = 12 patients of both sexes was immediately placed in ice-cold sucrose aCSF (0°C±1°C) to ensure maximum viability.

### 3.2 Electrophysiological Recordings

Resected tissue was kept in ice cold sucrose until sliced using a submerged vibratome (Leica VT1200S, Leica Systems, Germany). The tissue was then incubated in sucrose aCSF (30 mins, 37°C) and followed by room temp (24°C±1°C) for 30 mins before patch clamp experiments were carried out.

Electrophysiological experiments were carried out at room temperature (24°C±1°C) in aCSF solution. Slices were visualized with an upright microscope (Olympus BX61WI, Olympus Corporation, Japan) with infrared differential interference (IR-DIC) optics. Whole cell recordings were obtained from cortical pyramidal cells (identified by their shape and prominent apical dendrite), from cortical layer III-IV. Recording pipettes were filled with an internal solution containing the following (in mM:). All substances and drugs were obtained from Sigma-Aldrich or Tocris Bioscience. Recordings were made using a Multiclamp 700B (Molecular Devices LLC, U.S.A.) amplifier and digitized using a CED 1401 MKII (Cambridge Electronic Design Ltd., U.K.) at 20 KHz. Signals were Bessel filtered at 8 KHz and acquired using a custom script in IGOR Pro (Wavemetrics Inc, U.S.A.). Series resistances were carefully monitored and the recordings were discarded if varied by more than 50%. The recorded traces were analyzed using Clampfit 10 (Molecular Devices LLC, U.S.A.). Action potentials in pyramidal cells were induced by stepwise depolarization from 0-0.5 nA, with .025 nA steps. Recordings were made once, after a 10-minute equilibration after the whole cell configuration was achieved.

Once the whole cell configuration was achieved and after an equilibration phase of 10 mins, recordings were performed with depolarization of 25 pA stepwise.

#### 5.3 Differences in Membrane resistance and Total conductance

The *input resistance* of a neuron reflects the extent to which membrane channels are open. A *low* resistance (high conductance) implies *open* channels, while *high* resistance implies *closed* channels. In practice, to measure input resistance, packets of charges (current pulses) are injected through a microelectrode, and the potential that results is measured. If the injected charges produce a big change in potential, few charges have leaked out across the membrane and the input resistance is high. If the injected charges produce a small change in potential, charges have leaked across the membrane and the input resistance is low (open channels have let the charges escape). Measuring input resistance while applying a transmitter shows whether the transmitter opens or closes channels. The membrane resistance R_m_ is calculated as below

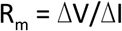

where ΔV is the difference in the V_m_ between two different current injections and ΔI is the current step (25 pA). Current injection steps from 0 pA to 200 pA were used to plot the IV curve and for estimating the membrane resistance R_m_. The membrane resistance can be used to estimate the number of channels that are present in the cell. non-epileptic cells have a mean resistance of 78.87±10.36 MΩ, while epileptic cells have a mean membrane resistance of 53.13±12.17 MΩ. There is a significant difference between the two, with a p value of0.0002, calculated using the *unequal variances t-test*. The conductance can be estimated as the inverse of the resistance. The conductance can also be estimated by the following

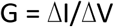

The mean conductance of non-epileptic cells is 12.86±1.576 and that of epileptic cells is 19.72 ±4.545. There is a significant difference between the two, given by a p=0.0017, calculated using the *unequal variances t-test*.

#### 3.3 In silico Modelling

A morphological and biophysically accurate model of human pyramidal neurons is currently not available. However, in this work we were not interested in modeling human cells but in understanding the possible changes underlying the differences in the dynamical properties between neurons from non-epileptic and epileptic tissue. For this reason, we used a highly validated computational model of a rodent’s pyramidal neuron, based on a set of channel kinetics previously used in other works (Ascoli et al., 2010; Migliore and Migliore, 2012) and a full 3D morphology. Simulations were carried out using the NEURON environment (v7.5) (Hines and Carnevale, 1997). As control condition, we started from a data-driven optimized model of a morphologically and biophysically detailed rat hippocampal CA1 pyramidal neuron, without attempting any tuning to obtain a quantitative match between experimental and modeling traces. The model has been obtained using one of the workflows available on Brain Simulation Platform of the Human Brain Project (BSP,https://www.humanbrainproject.eu/en/brain-simulation/). Complete model files are available after publication on the BSP (file CA1_pyr_cACpyr_dend-050921AM2_axon-051208AM2_20170516112638) and on the public Model DB database (https://senselab.med.yale.edu/modeldb/, acc.n.231106).

#### 3.4 Data Analysis

Action potential analysis was carried out in Clampfit 10 (Molecular Devices Inc., USA) and further analysis was carried out in Graphpad Prism. All comparisons were carried out using the non-parametric Mann-Whitney test.

## 4. Author Contributions

K.J. and V.M.R conducted experimental and theoretical work respectively. J.Z. provided access to resected human tissue otherwise discarded. V.M.R., R.M. and M.M. conducted the computational modelling. U.G.H. and M.M. supervised the experimental and theoretical work. K.J., V.M.R., T.J.F., R.M., M.M. and U.G.H. wrote and reviewed the manuscript.

## 5. Acknowledgements

The authors would like to acknowledge the surgical team of the Department of Neurosurgery - Medical Center University of Freiburg and C. Donkels for timely access to resected tissue. The authors would like to thank C. Bernard for helpful discussions.

